# Improved cohesin HiChIP protocol and bioinformatic analysis for robust detection of chromatin loops and stripes

**DOI:** 10.1101/2024.05.16.594268

**Authors:** Karolina Jodkowska, Zofia Parteka-Tojek, Abhishek Agarwal, Michał Denkiewicz, Sevastianos Korsak, Mateusz Chiliński, Krzysztof Banecki, Dariusz Plewczynski

## Abstract

Chromosome Conformation Capture (3C) methods, including Hi-C (a high-throughput variation of 3C), detect pairwise interactions between DNA regions, enabling the reconstruction of chromatin architecture in the nucleus. HiChIP is a modification of the Hi-C experiment, which includes a chromatin immunoprecipitation step (ChIP), allowing genome-wide identification of chromatin contacts mediated by a protein of interest. In mammalian cells, cohesin protein complex is one of the major players in the establishment of chromatin loops. We present an improved cohesin HiChIP experimental protocol. Using comprehensive bioinformatic analysis, we show that performing cohesin HiChIP with two cross-linking agents (formaldehyde [FA] and EGS) instead of the typically used FA alone, results in a substantially better signal-to-noise ratio, higher ChIP efficiency and improved detection of chromatin loops and architectural stripes. Additionally, we propose an automated pipeline called nf-HiChIP (https://github.com/SFGLab/hichip-nf-pipeline) for processing HiChIP samples starting from raw sequencing reads data and ending with a set of significant chromatin interactions (loops), which allows efficient and timely analysis of multiple samples in parallel, without the need of additional ChIP-seq experiments. Finally, using novel approaches for biophysical modelling and stripe calling we generate accurate loop extrusion polymer models for a region of interest and a detailed picture of architectural stripes, respectively.

## Introduction

Genome folding is a complex process that efficiently packs roughly two-metre-long human DNA into a cell nucleus that is only a few micrometres in diameter. Importantly, it does not serve only structural purposes, but must be tightly coordinated with fundamental processes occurring in the nucleus, such as DNA replication, transcription, or DNA repair. Despite significant progress in 3D genomics in recent years, our understanding of the relationship between genome structure and function is still largely elusive and requires further investigation. There is a need for improved methods to detect chromatin contacts at different resolutions and for bioinformatic pipelines that comprehensively and efficiently analyse complex experimental data.

Studies of DNA structure in the nucleus using 3C-based technologies and microscopic imaging methods have shown that chromatin is folded at different levels of organisation in the nucleus. At the lower scale, interphase chromosomes occupy discrete regions within the nucleus called chromosome territories (1, 2). They are partitioned into alternating multi-megabase scale regions called compartments of two types, A and B, corresponding to euchromatin and heterochromatin, respectively (3). Furthermore, Topologically Associated Domains (TADs) can be detected within the compartments. TADs are sub-megabase regions that have stronger contacts within themselves than with other regions (4–6). At a finer scale, the 10 chromatin fibre folds into loops which connect elements that may be distant in the linear genome (such as promoters and enhancers). A subset of loops has been shown to act as boundaries of TADs (6). Importantly, both TADs and loops were shown to be stable across different cell lines and even between different species (4, 6, 7).

In mammalian cells, the major factors involved in TAD and loop formation are the evolutionarily conserved proteins cohesin and CCTC-binding factor (CTCF). Cohesin is a ring-shaped protein complex composed of four subunits: two structural maintenance of chromosomes (SMC) proteins, SMC1 and SMC3, and two non-SMC proteins: RAD21 and stromal antigen (SA) that in vertebrates comes in two versions STAG1 (SA1) or STAG2 (SA2, reviewed in (8)). Cohesin association with the DNA is dynamic: it encircles the DNA upon loading (9) and is able to slide along the DNA (10, 11). The sites of cohesin loading differ from the sites of its final genomic localisation (12). The transcription process was shown to translocate cohesin along the DNA over long distances (11, 13). CTCF is a sequence-specific DNA binding protein that contains a highly conserved 11 zinc finger domain. CTCF, together with cohesin, is enriched at TAD boundaries (4, 5) and at the majority of loop anchors (3, 6). In mammalian cells, cohesin colocalises with CTCF in most of its binding sites (14, 15).

It was proposed (16–18) that TADs and loops are formed by a loop extrusion process, in which cohesin uses its ATP-dependent motor function to move along the DNA, causing active loop enlargement until it encounters a physical barrier, such as the CTCF protein, or until it dissociates from chromatin (19–21). By stopping the extrusion process, CTCF acts as a chromatin loop anchor. It restricts the contacts occurring within the TADs and prevents them from crossing their borders. While CTCF-mediated interactions are conserved to a great extent across cell types and during differentiation, CTCF-independent interactions have a more dynamic nature; they are cell type-specific, associated with gene expression, and often reside at enhancers (22, 23).

3C techniques, including Hi-C, detect interactions between different DNA regions, enabling the reconstruction of the chromatin architecture in the nucleus (reviewed in (24). A standard Hi-C procedure involves chromatin structure fixation, restriction enzyme digestion, end repair, and ligation of fragments localised close to each other in the nuclear space. The DNA is then fragmented to generate linear chimeric fragments, which is followed by genomic library preparation, high-throughput sequencing, and bioinformatic analysis. Modifications of Hi-C experiments, such as HiChIP (25), PLAC-seq (26), similarly introduced in 2009 ChIA-PET methodology (27) use a chromatin immunoprecipitation (ChIP) step, for genome-wide detection of interactions mediated by a specific protein. In the HiChIP approach, ChIP with the antibody of interest is performed after the Hi-C ligation step (**Figure 1A**). An important advantage of the HiChIP approach over the standard Hi-C is that a higher resolution can be achieved with a lower sequencing depth.

**Figure 1.**
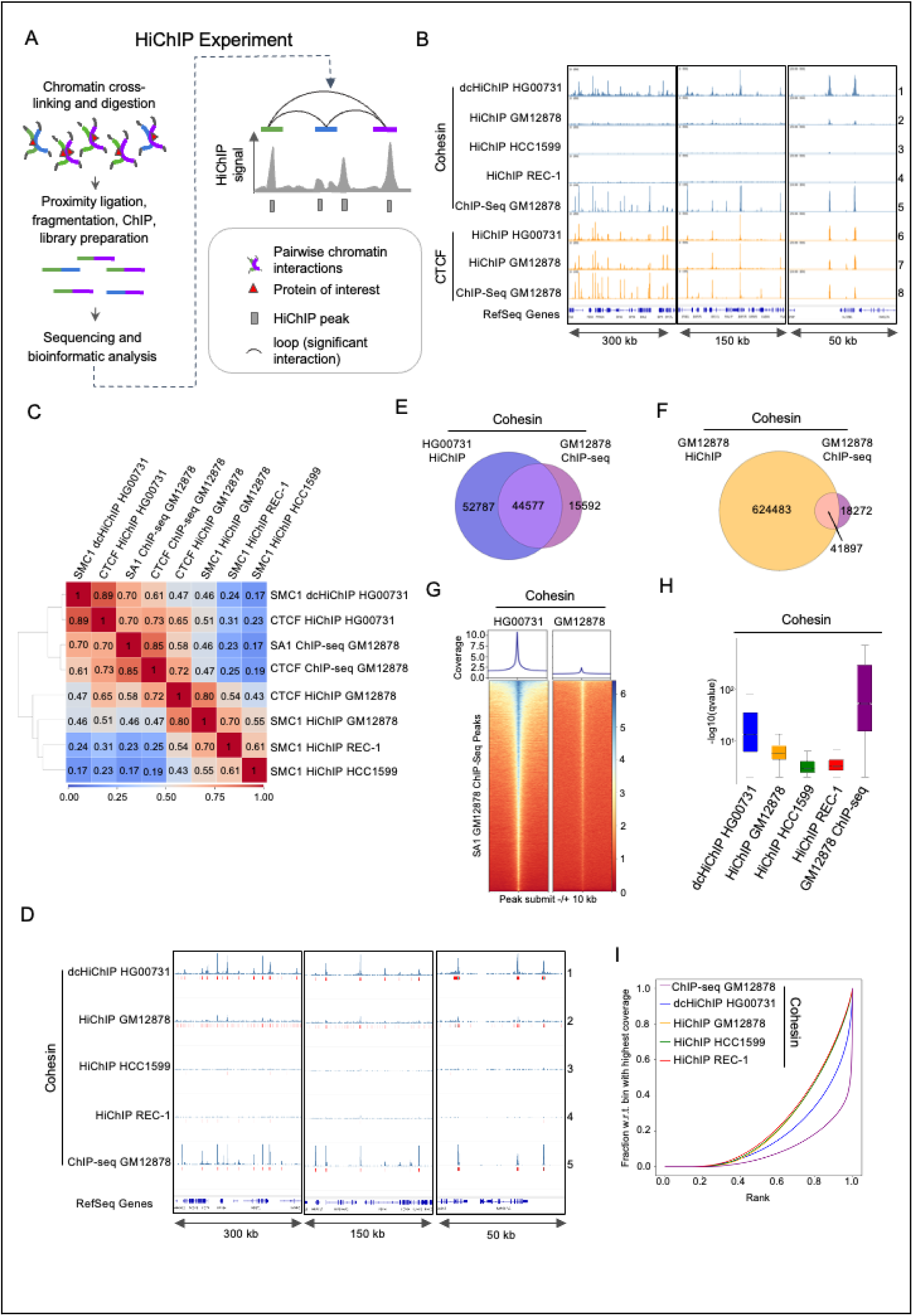
FA-EGS crosslinking improves the quality of the cohesin HiChIP experiment. **A.** Schematic of the HiChIP experiment and its data representation. **B.** IGV browser coverage tracks of the HiChiP and ChIP-seq experiments analysed in this study. The following exemple genomic regions are shown: (1) 300 kb window – chr19:45,417,401-45,717,713; (2) 150 kb window – chr19:43,508,033-43,658,922; (3) 50 kb window – chr17:49,677,416-49,727,883. **C.** Heatmap showing Pearson correlation coefficient between the samples processed in this study. **D.** IGV browser coverage tracks and called peaks for the indicated cohesin HiChIP (SMC1) and ChIP-seq (SA1) experiments. The following exemple genomic regions are shown: (1) 300 kb window – chr5:150,332,698-150,635,698; (2) 150 kb window – chr7:55,939,000-56,089,000; (3) 50 kb window – chr17:40,160,000-40,210,000. **E.** Venn diagrams showing common peaks between SMC1 HiChIP HG00731 and SA1 ChIP-seq GM12878. **F.** Venn diagrams showing common peaks between SMC1 HiChIP GM12878 and SA1 ChIP-seq GM12878**. G.** Heatmap showing the distribution of SMC1 FA-EGS HiChIP HG00731 reads and SMC1 HiChIP GM12878 reads within a 20 kb window centred on the cohesin (SA1) ChIP-seq GM128787 peaks. **H.** Distribution of –log(10)qvalue of peaks called by MACS3 in the indicated cohesin HiChIP and ChIP-seq samples. Red line indicates the median. **I.** Fingerprint analysis plot (45) showing the comparison of ChIP enrichment in the indicated cohesin HiChIP and ChIP-seq samples.

HiChIP experiments require a extensive computational analysis to determine chromatin structure features such as: (a) HiChIP peaks detected from the signal of aligned sequencing reads, (b) chromatin interaction matrices showing all contacts detected in a given sample, (c) significant DNA interactions (loops) corresponding to the most frequent chromatin interactions present in the cell population, and finally (d) architectural stripes appearing on the contact matrices as horizontal or vertical lines (17). Stripes are postulated to be the effect of loop extrusion, where one loop anchor is held in place, and becomes the stripe anchor, and the other loop anchor moves through the stripe domain until the loop is fully extruded (28, 29). During the extrusion process promoters can be brought into contact with long clusters of enhancers (17).

Dedicated methods have been developed to analyse specific 3C experiments, such as Juicer (30) for Hi-C, MAPS (31) for HiChIP and PLAC-seq data, and ChIA-PIPE (32) for ChIA-PET. In this study, we tested all three pipelines to analyse and compare our improved HiChIP protocol with publicly available HiChIP datasets. We also used two different stripe-calling algorithms: Stripenn (33), a state-of-the art tool operating on contact matrices, and the novel tool called gStripe, which is specifically designed to analyse sparse data using graph methods (see Methods for a description).

We observed that the quality of the cohesin HiChIP data is significantly lower compared to the CTCF HiChIP. This could be due to the more dynamic, mobile nature of the cohesin complex compared to CTCF, as discussed above. In this study, we present a modified version of the cohesin HiChIP protocol that significantly improves signal-to-noise ratio and loop detection efficiency. Importantly, we propose an automated pipeline “nf-HiChIP”, based on the Nextflow system (34), for efficient and timely analysis of HiChIP samples starting from raw sequencing reads and ending with a set of significant chromatin interactions (loops). This pipeline can be easily deployed on a local workstation, a high performance computing system, or cloud-based computing environments. Finally, using high-quality cohesin HiChIP data obtained with improved protocol and a novel biophysical modelling approach for loop extrusion (35), we simulated dynamic 3D models of loop extrusion for a region of interest.

## Results

### Standard cohesin HiChIP protocol yields results with low signal-to-noise ratio

This study focuses primarily on human lymphoblastoid cell lines (LCLs) from the 1000 Genome Project (36), a widely used biological model for genetic and genomic research. We observed that the publicly available cohesin (SMC1) HiChIP for human GM12878 LCL (25) yields a much lower signal-to-noise ratio than the CTCF HiChIP performed with HC 3C protocol for the same cell line (37), and than the CTCF HiChIP for HG00731 LCL performed in our laboratory (**Figure 1B**, lanes 2, 6, 7; **Supplementary Figure 1B**). A low signal-to-noise ratio was also observed in publicly available cohesin (SMC1) HiChIP experiments performed in the human lymphoblast cell line REC-1 and the human epithelial cell line HCC1599 ((38), **Figure 1B**, lanes 3 and 4). Visual comparison of the cohesin HiChIP and cohesin (SA1) ChIP-seq (39) signal in GM12878 indicates a limited enrichment of sequencing reads around cohesin ChIP-seq peaks in HiChiP experiments (**Figure 1B**, lanes 2 and 5).

To improve the quality of the HiChIP experiment, we tested a modified HiChIP protocol (**Supplementary Figure 1A**) that includes an additional formaldehyde (FA), cross-linking agent – EGS, suggested to improve signal-to-noise ratio in ChIP experiments (40, 41). The dual cross-linking protocol was previously used for the ChIA-PET experiments (42, 43), and it was also recently shown to improve the quality of the Hi-C experiment (44). For clarity, we have referred to the modified HiChIP protocol as FA-EGS HiChIP or dual cross-linking HiChIP (dcHiChIP) to distinguish it from the standard protocol. The summary of the HiChIP experiments used in this study is provided in **Supplementary Table 1**.

### Dual cross-linking protocol improves cohesin ChIP efficiency and enables successful detection of cohesin HiChIP peaks

We performed cohesin FA-EGS HiChIP experiments for the HG00731 LCL in two biological replicates. By visual inspection in the genome browser and using the Pearson correlation coefficient, we confirmed a high reproducibility between two replicates for the following experiments: (1) cohesin FA-EGS HiChIP for HG00731, (2) CTCF HiChIP for HG00731, (3) cohesin (25) and (4) CTCF (37) HiChIP for GM12878 (**Supplementary Figure 1B and C**). Therefore, we pulled both replicates for further analysis. Cohesin HiChIP experiments in REC-1 and HCC1599 cell lines (38) were performed in one replicate and, as such, were used in the following analysis.

To first assess the quality of the dual cross-linking approach, we first compared cohesin FA-EGS HiChIP with standard cohesin HiChIP experiments. For reference, we also used cohesin and CTCF ChIP-seq experiments for GM12878. Unfortunately, ChIP-seq experiments for HG00731 LCL were not available, and therefore, we could not include them in the analysis. Visual inspection in the genome browser indicates a higher signal-to-noise ratio in the sequencing coverage track (**Figure 1B, Supplementary Figure 1B**) of cohesin FA-EGS HiChIP compared to FA HiChIP experiments and a comparable signal-to-noise ratio to CTCF HiChIP experiments. Next, we examined the similarity between the analysed experiments using the Pearson correlation coefficient (**Figure 1C**). As expected, the highest correlation is observed between cohesin and CTCF HiChIP or cohesin and CTCF ChIP-seq experiments performed for the same cell line (GM12878 and HG00731). Interestingly, cohesin FA-EGS HiChIP (HG00731) showed a better correlation with GM12878 cohesin ChIP-seq (r = 0.70) than with GM12878 cohesin HiChIP (r = 0.46), despite the fact that different LCLs were compared. This is consistent with what was suggested by visual inspection of the tracks (**Figures 1B, lanes 1, 2 and 5**). A moderate correlation between cohesin dcHiChIP (HG00731) and cohesin HiCHIP (GM12878, r = 0,46) and poor correlation with cohesin HiChIP performed for REC-1 and HCC1599 (0.24 and 0.14 respectively) was also observed (**Figure 1C**). We conclude that dual cross-linking enhances the quality of the cohesin HiChIP signal and that the limited correlation between cohesin dcHiChIP and standard HiChIP could be caused by the lower signal-to-noise ratio in the former type of experiment.

To better understand the difference between dcHiChIP and standard cohesin HiChIP we performed peak calling using the MACS3 algorithm. Importantly, in the HiChIP experiment, peaks may correspond to regions that are bound by a target protein or to regions that are in contact with the target protein binding sites. Therefore, a higher number of peaks can be expected for the HiChIP experiment compared to the ChIP-seq experiment. As no input sample is generated in the HiChIP experiment, the MACS3 function for calculating the local background of the HiChIP sample (λ_local_) was used for peak enrichment analysis. The high overlap between peaks, called in separate replicates further confirmed the reproducibility of the HiChIP experiments (**Supplementary Figure 2A**). Over 600000 and nearly 100000 peaks were detected for cohesin HiChIP (GM12878) and FA-EGS HiChIP (HG00731), respectively. For the cohesin HiChIP experiment performed in REC-1 and HCC1599 a much lower number of peaks was found (34379 and 21281, respectively). For comparison, MACS3 algorithm detected for cohesin and CTCF ChIP-seqs in GM12878 LCL approximately 60000 and 67000 peaks, respectively **(Supplementary Table 2)**. We were surprised by the high number of peaks detected for cohesin HiChIP in GM12878 (25) despite a rather low signal-to-noise ratio. Indeed, the genome browser view shows that some of the peaks are located in the regions that are not very distinct from the noise (**Figure 1D, lane 2**). This was not the case for the FA-EGS HiChIP protocol, where peaks are distinct from the background noise (**Figure 1D, lane 1**). We hypothesise that the combination of the low signal-to-noise ratio of the data and the lack of input control may be a reason why the MACS algorithm detected so many peaks in this particular HiChIP experiment.

Importantly, 74% of the cohesin ChIP-seq peaks overlapped with the cohesin FA-EGS HiChIP peaks (**Figure 1E**). We consider this a significant overlap taking into account that the experiment was performed in different LCLs (GM12878 vs. HG00731) and using different experimental protocols and antibodies targeting different cohesin subunits (SA1 vs. SMC1). For comparison, the overlap between cohesin HiChIP and ChIP-seq peaks for the GM12878 cell line was slightly lower (70%), although data were derived from the same cell line and that the number of HiChIP peaks was much higher **(Figure 1F)**. From this analysis we conclude that SMC1 dcHiChIP accurately detects cohesin binding sites. Moreover this analysis indicates that when HiChIP data have a high signal-to-noise ratio, as is the case for the cohesin FA-EGS HiChIP, the lack of input control is not an obstacle to reliable HiChIP peak detection by the MACS peak calling algorithm.

To verify the influence of dual cross-linking on ChIP, we analysed the enrichment of cohesin (SMC1) GM12878 HiChIP and HG00731 FA-EGS HiChIP reads around cohesin (SA1) GM12878 ChIP-seq peaks (39). Strikingly, the enrichment of cohesin HG00731 FA-EGS HiChIP reads around GM12878 cohesin ChIP-seq peaks was substantially higher compared to GM12878 cohesin HiChIP **(Figure 1G).** Furthermore, the significance levels of the set of peaks measured using the q-value parameter (MACS3) showed the highest score for FA-EGS HiChIP compared to other HiChIP samples (**Figure 1H**). We also performed Fingerprint analysis (45, 46). According to this tool the better the quality of the ChIP sample is, the smaller the area under the curve (AUC) and the higher the elbow point it presents. As a reference we used the cohesin ChIP-seq experiment which, as expected, showed the highest quality. The FA-EGS HiChIP showed the best quality among the cohesin HiChIP examined (**Figure 1I).** These results indicate that the efficiency of the ChIP step in the cohesin FA HiChIP protocol is limited and that the dual cross-linking step improves it. Our results confirm that HiChIP provides reliable information about cohesin binding to DNA, as previously reported (25). We conclude that FA-EGS cross-linking improves the signal-to-noise ratio, increases the efficiency of ChIP step in cohesin HiChIP experiment and enables reliable detection of HiChIP peaks.

### Dual cross-linking HiChIP protocol improves detection of cohesin-mediated loops

In order to detect loops (significant interactions) connecting two distant genomic fragments (anchors), we used three independent algorithms: 1) nf-HiChIP and 2) ChIA-PIPE (32), tools specifically designed to process experiments targeting protein-mediated interactions (such as HiChIP, PLAC-seq or ChIA-PET) as well as, 3) Juicer (30), an algorithm originally developed for Hi-C samples. The first algorithm, nf-HiChIP, is based on MAPS (31) – an algorithm used for loop calling – which takes as input mapped, paired and filtered reads, as well as the binding site information of the protein of interest (usually ChIP-seq experiment). The nf-HiChIP extends the algorithm by allowing the input to be raw sequences, calculating all the necessary preprocessing steps automatically. Identified interactions have at least one anchor that is enriched in a given protein factor. Juicer, on the other hand, generates interaction matrices (contact maps) which are then processed by a loop-calling algorithm (HiCCUPS). It identifies interactions by detecting groups of pixels representing the most frequent interactions, visible as “dots” on the contact maps. Therefore, loop detection by Juicer is independent of ChIP-seq peaks. ChIA-PIPE takes paired-end reads and provides clusters of the paired-end tags (PETs). It then merges overlapping PETs to determine the PET counts of potential chromatin contacts (or “looping”) frequency between two anchor loci involved in chromatin interaction to estimate the strength of the loop.

We developed a multipurpose, Nextflow-based (34) pipeline called nf-HiChIP (https://github.com/SFGLab/nf-hichip) designed for NGS sequencing data analysis which performs (i) peak calling for ChIP-Seq and HiChIP data and (ii) loop calling (i.e., identification of significant contacts) from HiChIP data **(Supplementary Figure 3**). The nf-HiChIP pipeline is compatible with multiple samples and replicates, streamlining the analysis of large datasets through automation. Implemented in Python3 (47) and using Nextflow, the pipeline excels at managing complex data workflows. Nextflow facilitates task scheduling and dependency resolution while providing robust error handling mechanisms – essential for large-scale analyses. The pipeline is structured into well-defined tasks, allowing for efficient error correction and resumption of the process from the last successfully completed task, eliminating the need to restart the analysis from scratch. The use of the pipeline is straightforward – installing can be done using 3 commands as shown in Nextflow documentation, and running HiChIP pipeline requires a sample design file. The rest of the workflow is fully automated, so the only input required is a .csv file describing the experiment and fastq files.

In our pipeline, in the loop calling step, we decided to use HiChIP peaks instead of ChIP-seq peaks. We reasoned that this solution would be particularly advantageous when the ChIP-seq data is not available for the cell line and/or antibody of interest, or when the available ChIP-seq data is of poor quality. To test whether this approach reliably detects chromatin interactions, we analysed the CTCF HiChIP (GM12878) sample, which has a high signal-to-noise ratio and for which CTCF ChIP-seq peaks are publicly available. We compared loops detected by the nf-HiChIP algorithm with CTCF ChIP-seq or CTCF HiChIP peaks and found that the overlap reached 98% (**Supplementary Table 3**). Additionally, we tested the similarity between nf-HiChIP loops (peak-dependent algorithm) and HiCCUPS loops, which are independent of peaks. For the HiCHIP samples analysed in this study, we have found that approximately 75% to 90% of nf-HiChIP loops were common with HiCCUPS loops (**Supplementary Table 4)** further validating the reliability of loops detected by nf-HiChIP using HiChIP peaks instead of ChIP-seq peaks. This confirms that when a HiChIP sample has a high signal-to-noise ratio, loop-calling analysis with nf-HiChIP can be performed using HiChIP peaks from the same sample instead of using ChIP-seq peaks from additional experiments.

Visual inspection in the IGV and Juicebox genome browsers shows that all three algorithms detected more chromatin interactions in the FA-EGS HiChIP sample, compared to the other cohesin HiChIP samples analysed (**Figure 2A-B, Supplementary Figures 4A-B and 5A-B**). Indeed, in cohesin dcHiChIP (HG00731), the number of detected loops was at least twice as high as in the other cohesin HiChIP experiments analysed and was 80487, 43148 and 179516 for nf-HiChIP, HiCCUPS and ChIA-PIPE, respectively (**Supplementary Table 5**). The enrichment score of detected loops was calculated using aggregate peak analysis (APA, (6)) with nf-HiChIP, Juicer, and ChIA-PIPE loops based on interaction maps generated by Juicer software. In cohesin FA-EGS HiChIP, the aggregate loop strength was significantly better as compared to the remaining cohesin HiChIP experiments. It was comparable (in some cases even better) to the APA scores obtained for CTCF HiChIP experiments (**Figure 2C, Supplementary Figure 4C and 5C**). Therefore, in the cohesin HiChIP experiment, dual cross-linking improves loop detection in terms of loop number and strength in comparison to the FA cross-linking.

**Figure 2.**
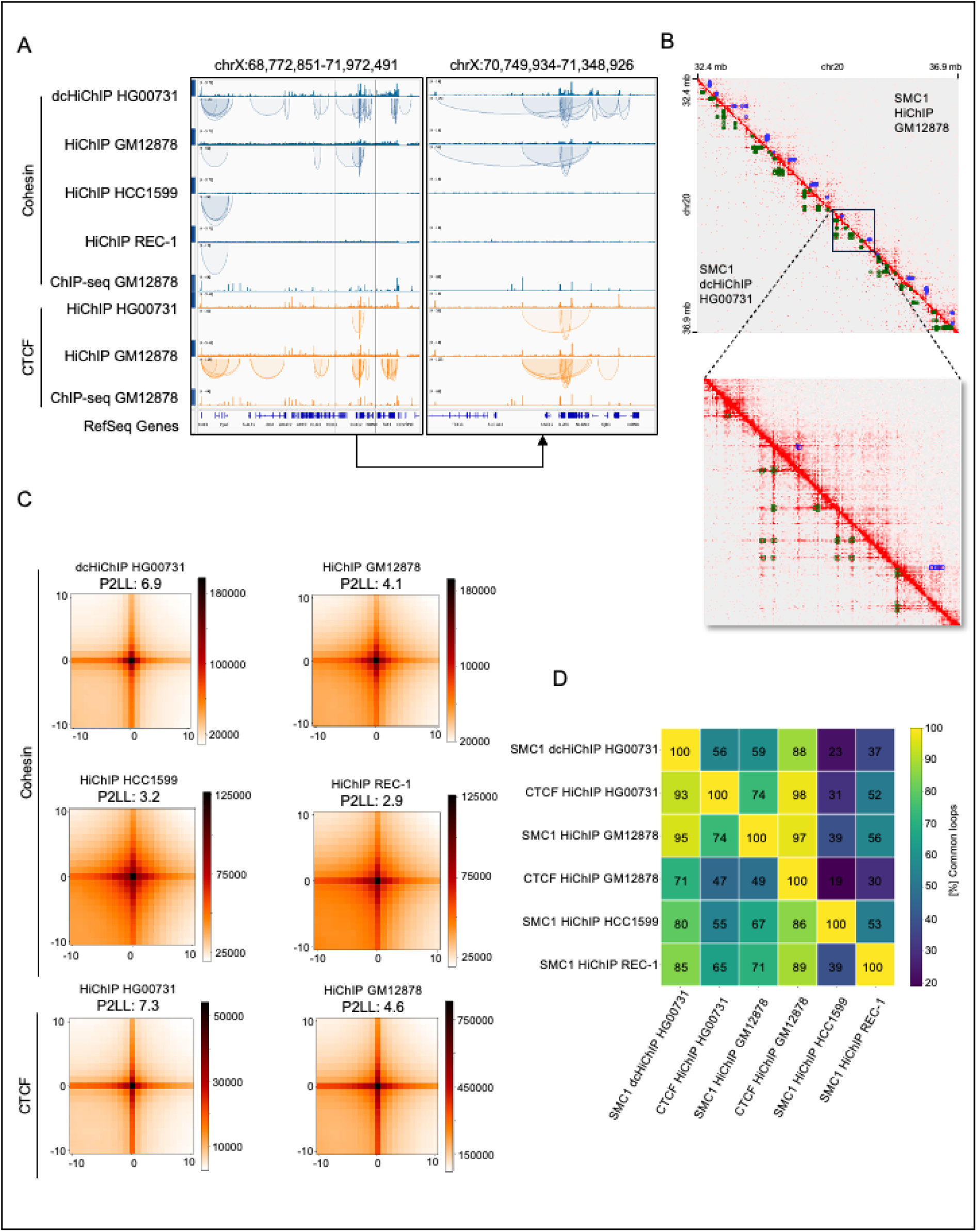
FA-EGS crosslinking HiChIP protocol improves cohesin-mediated loop detection. **A**. IGV browser coverage tracks and nf-HiChIP loops of the indicated HiChIP samples and read coverage tracks for SA1 and CTCF GM12878 ChIP-seq experiments. **B.** Juicebox interaction maps at 5 kb resolution of the exemple region for the indicated cohesin HiChIP samples. Loops (nf-HiChIP) are shown as green (SMC1 HiChIP HG00731) or blue (SMC1 HiChIP GM12878) rectangles. **C**. APA analysis performed using loops identified by nf-HiChIP for the indicated cohesin and CTCF HiChIP samples. The APA score P2LL (peak to left lower corner) is the ratio of the central bin to the average of the lower left corner and indicates the strength of the loop. **D.** Overlap between loops called by nf-HiChIP in the indicated datasets. A single cell in the matrix represents the percentage of loops in the row sample that overlap with one or more loops from the column sample. For example, 93% of the loops from the HG00731 CTFC (row 2) and 95% of the loops from the GM12878 SMC1 (row 3) were present in HG00731 SMC1 (column 1). Note, that the matrix is not symmetric, as one-to-many overlaps are possible.

Visual examination of the loops in the genome browsers identified from nf-HiChIP and HICCUPS with the fixed anchor size of 5kb suggested that, despite the differences in the number of contacts detected (**Supplementary Table 5**), there is a high degree of concordance in contact localisation between cohesin FA-EGS HiChIP and other cohesin HiChIPs analysed (**Figure 2A-B, Supplementary Figure 4A-B**). Whereas in ChIP-PIPE the identified loops are with flexible anchor size and number of cohesin FA-EGS HiChIP loops was quite high as compared to other cohesin HiChIPs (**Supplementary Table 5**) therefore visual examination of the ChIA-PIPE loops are more dense (**Supplementary Figure 5A-B**). Indeed, the loop overlap between cohesin FA-EGS HiChIP (HG00731) and cohesin HiChIP obtained for GM12878, REC-1 and HCC1599 was 95%, 80% and 85% (nf-HiChIP) respectively (**Figure 2D**). We observed similar concordance in HiCCUPS interactions (**Supplementary Figure 4D**), where within the interaction identified from ChIA-PIPE we identified similar behaviour but with slightly lower overlap percentage with the short and short read sequencing samples HCC1599 and REC-1 (**Supplementary Figure 5D**). This confirms the reliability of the improved HiChIP protocol and indicates that it allows for the generation of more complex maps of cohesin-mediated chromatin contacts than the standard one. The comparison between cohesin FA-EGS HiChIP and FA HiChIP shows that the general pattern of the sequencing signal on the interaction matrices is similar between experiments, but standard protocol produces a more diffused signal (**Figure 2B** and **Supplementary Figure 4B and 5B**).

Of note, not all peaks called from the raw sequencing data will be involved in chromatin interactions. Analysis of the proportion of HiChIP peaks that localise inside the loop anchors (3D peaks) detected by nf-HiChIP revealed that this percentage was the highest for cohesin FA-EGS HiChIP (51%), followed by cohesin HiChIP in HCC1599 and CTCF HiChIP in GM12878 (39% and 28% respectively). The lowest proportion of the 3D peaks within HiChIP anchors was observed for both CTCF and cohesin HiChIP peaks in GM12878 LCL and cohesin HiChIP in REC-1 which are 15%, 11% and 17% respectively (**Supplementary Table 2**). Interestingly, 3D peaks present higher cohesin or CTCF enrichment than peaks that are not involved in chromatin interactions (**Supplementary Figure 7A**) suggesting their greater biological relevance. As expected, the analysis of CTCF motif orientation within loop anchors revealed that CTCF motifs are positioned predominantly in convergent orientation for all the HiChIP experiments analysed (**Supplementary Figure 7B**).

### Dual cross-linking HiChIP protocol reveals architectural stripes in more detail

To assess the quality of the FA-EGS HiChIP data with respect to the detection of structures above the loop level, we perform the calling of architectural stripes. Stripe calling was performed on the cohesin HiChIP datasets using the Stripenn tool (33), which is based on image processing, and a novel method recently developed in our laboratory called gStripe, based on graph analysis (see Materials and Methods for details). Stripenn uses as input the heatmaps generated by Juicer, while gStripe operates on loops obtained in previous steps from either nf-HiChIP, HiCCUPS or ChIA-PIPE tools.

First, we compared how many stripes could be extracted from each dataset. The number of stripes called by Stripenn in the cohesin FA-EGS HiChIP sample (HG00731) is 8% lower than in the cohesin HiChIP (GM12878) sample (3603 vs 3908), but more than two and three times higher in comparison to REC-1 and HCC1599 cohesin HiChIP samples, respectively (**Figure 3A, Supplementary Table 6**). On the other hand, gStripe consistently detected many more stripes in cohesin datasets derived from FA-EGS HiChIP, regardless of the loop-calling tool used. In particular, gStripe detected two to four times more stripes for cohesin FA-EGS HiChIP compared to standard cohesin HiChIP (GM12878) using nf-HiChIP, HiCCUPS and ChIA-PIPE input loops.

**Figure 3.**
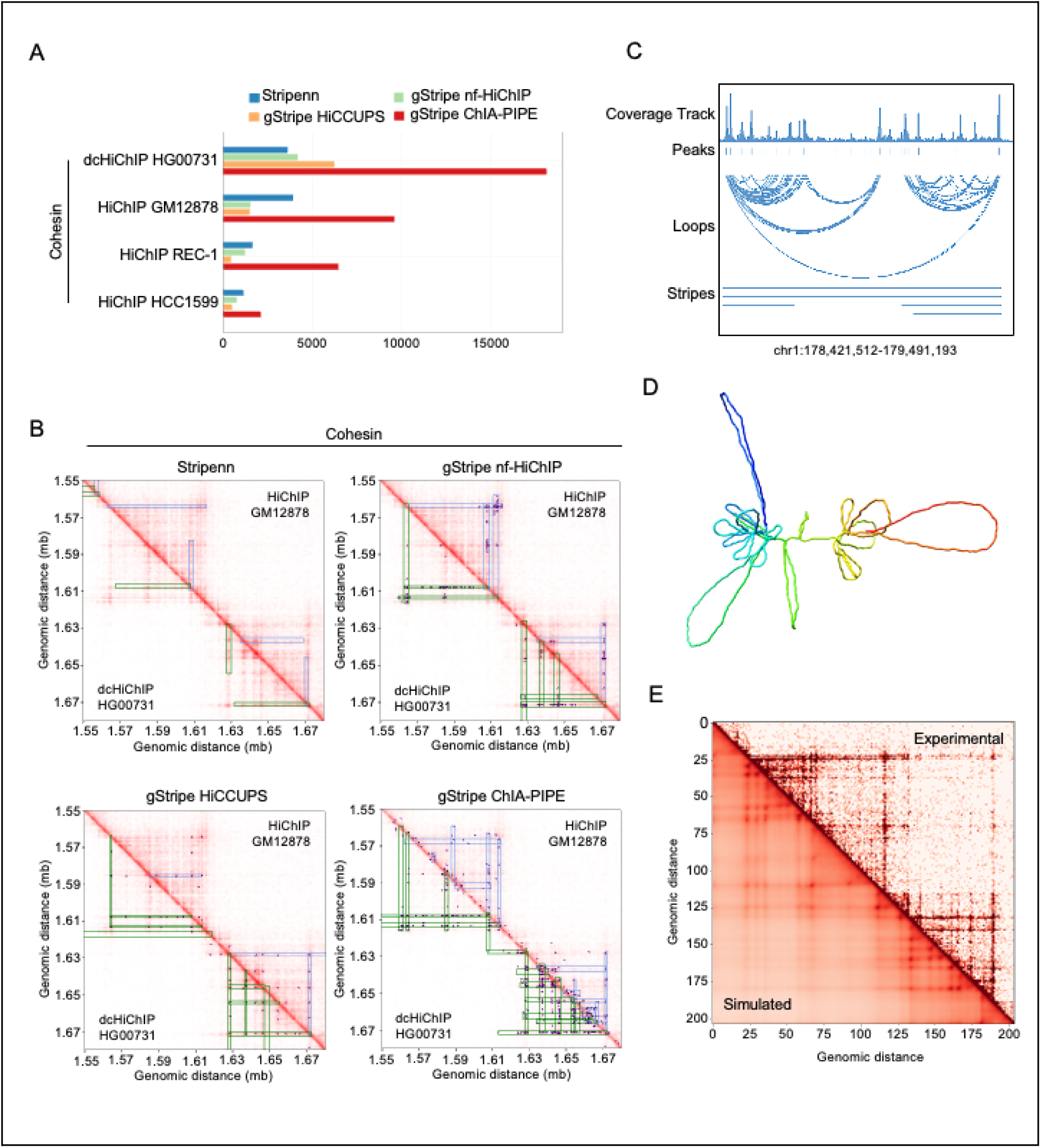
FA-EGS crosslinking HiChIP protocol improves stripe detection and enables the building of accurate loop extrusion models. **A**. Number of stripes called by Stripenn and gStripe for the cohesin HiChIP datasets analysed in this study. gStripe results were calculated using nf-HiChIP, HiCCUPS or ChIA-PIPE loops as an input. **B.** Juicebox interaction maps at 5 kb resolution and the stripes of the exemplary region (chr6: 15,550,000-16,850,000) for the cohesin HiChIP samples. Lower (below diagonal) and upper (above diagonal) corners present SMC1 FA-EGS HiChIP (HG00731) and SMC1 HiChIP (GM12878), respectively. Stripes were called by the algorithms indicated above the maps. Each stripe set is shown on top of data used to generate it: the interaction heatmap and, in the case of gStripe, the nf-HiChIP, HiCCUPS or ChIA-PIPE loop set (blue dots). **C.** IGV browser view of the example region for SMC1 HG00731 FA-EGS HiChIP sample used for loop extrusion modelling. The first three tracks show the output of the nf-HiChIP pipeline: read coverage, peaks called by MACS3 and loops called by nf-HiChIP. The fourth track shows the stripes identified by gStripes (with the nf-HiChIP loop set). **D**. Final structure obtained after running LoopSage simulation for the region and sample indicated in panel C. **E.** Experimental (above diagonal) versus simulated(below diagonal) heatmaps (at the 5 kb resolution) for the region and data shown in panel C.

To get a more complete picture, we visually compared the stripe regions obtained from both protocols, and observed that often what appears as a smooth stripe signal in the standard HiChIP, often contains chains of distinct dots (loops) in FA-EGS HiChIP, (see e.g. the uppermost stripe in **Figure 3B** or **Supplementary Figure 8A**). Indeed, the Stripenn stripes in cohesin FA-EGS HiChIP (HG00731) contain 1.5, 1.8 and 2.6 times (for HiCCUPS, nf-HiChIP and ChIA-PIPE loops respectively) more loops per unit of area on the heatmap, than in cohesin FA HiChIP (GM12878, see **Supplementary Table 7**). As the image processing based Stripenn method expects a smooth signal, some of these stripes may be disturbed However, they can be recovered by gStripe. This is consistent with our preliminary observation that the FA-EGS HiChIP data appear to be more sensitive to changes in the parameters of Stripenn: within the same set of parameters of stripe calling, the ratio of the highest to lowest number of stripes called was 15.3 for HG00731 and only 2.5 for GM12878 (see **Supplementary Table 7** for the full summary).

Next, we investigated how the stripes are situated in relation to each other, by calculating the proportion of stripes located within the domain of another, larger stripe (see **Figure 3B** for an example). The stripe domain is defined as the range of coordinates from the stripe anchor to the end of the stripe. We found that a substantial proportion of the gStripe stripes are located within the domain of a larger stripe (between 57% when using nf-HiChIP loops, to up to 86% for ChIA-PIPE loops in cohesin FA-EGS HiChIP in HG00731 and between 38% and 70% for cohesin FA HiChIP in GM12878 respectively). We can therefore conclude that they often co-localise. On the other hand, the percentage of co-localised Stripenn stripes is lower: 21% for cohesin FA-EGS HiChIP (HG00731) and 31% for cohesin standard HiChIP (GM12878). In summary, the additional stripes detected by gStripe in cohesin dcHiChIP (HG00731) are smaller and more often co-localized within larger ones, especially when using a dense loop set such as the ChIA-PIPE, in comparison to standard cohesin HiChIP **(Figure 3B and Supplementary Figure 8A**).

Finally, to assess the reliability of the stripe detection, we check how many stripes from one dataset can be found in other sets. We look both at the overlap of individual stripe anchors and whole stripe domains. In the cohesin FA-EGS HiChIP (HG00731) Stripenn found approximately 49% of the stripes and 74% of the stripe domains detected in GM12878, 52% stripes and 78% domains from REC-1 HiChIP, and 43% stripes and 76% domains from HCC1599. For the gStripe algorithm, the degree of stripe overlap depends on the input loops set, but in general the overlap of stripes with the FA-EGS HiChIP (HG00731) was higher for cohesin HiChIP in GM12878 (e.g. 75% for nf-HiChIP loops) than for REC-1 and HCC1599 loops (e.g. 59% and 51% respectively, for nf-HiChIP). In the case of stripe domains, the overlap is very high: from 82% of HCC1599 domains recovered in HG00731 (using the HiCCUPS loop set) up to 98% of GM12878 domains (also for the HiCCUPS loops). The high overlap confirms the validity of the detected stripes.

Importantly, gStripe is able to recover a substantial number of GM12878 stripes (up to 88% when using HiCCUPS loop set and stripe domains (98%, also for the HiCCUPS loop set) Since gStripe detected more stripes in FA-EGS HiChIP (HG00731), we can say that in this data, the set of visible stripes and stripe domains is extended, as compared to GM12878. A complete set of overlap statistics between stripe anchors and stripe domains are shown in **Supplementary Figures 8B** and **Supplementary Figure 8C**, respectively. Taken together, our results suggest that the FA-EGS HiChIP experiment provides similar quality of detection for stronger, more uniform stripes using an established image-driven method, Stripenn, while having an advantage in revealing finer, co-located, stripes and uneven stripe structure when using a graph-based algorithm, gStripe.

### Dual cross-linking HiChIP protocol allows us to build accurate loop extrusion models

Finally, we aimed to generate biophysical models of the loop extrusion process for an exemplary region on chromosome 1 (**Figure 3C**), Usually, to simulate loop extrusion, stochastic models are used (16, 28, 48–50), where the cohesin proteins on the chromatin fibre perform one-dimensional random walk motion, and they may unload or load in different regions of chromatin stochastically. For the purpose of this work, we used our recently developed method, LoopSage (35), which combines stochastic simulation and molecular dynamics with OpenMM. In this model, cohesin rings play the role of loop extruders, and they follow a random walk diffusive motion, whereas CTCF acts as orientation-dependent barriers for its motion (18, 51). It also assumes that the position of CTCF proteins is more stable in comparison to cohesin complex (52, 53).

During the modelling process, first CTCF orientation within HiChIP loop anchors is determined. Next, LoopSage projects those sites in one dimension and constructs the binding energy of the system, which is composed of a leftward and a rightward part that encodes the orientation-specific barrier property of CTCF. By importing a HiChIP coverage track file, cohesin rings are distributed according to its preferential binding along the DNA. We run a simulation for SMC1 FA-EGS HiCHIP (HG00731). As the temperature decreases, we end up with a dynamical trajectory of 3D structures, where we assume that the final structure is the most representative one (**Figure 3D)**. We observe that the final polymer is condensed into two dominant clusters of loops, which correspond to the two squares in the experimental and reconstructed simulation heatmaps (**Figure 3E**). Comparison between the simulated heatmap, which is averaged over the ensembles of 3D structures and the experimental SMC1 dcHiChIP heatmap shows that loop and stripe patterns are well reconstructed (Pearson correlation between experimental and simulated map is 98.3%), indicating that our biophysical assumptions were valid.

## Discussion

In this study, using a comprehensive bioinformatics approach (summarised in the **Figure 4**), we show that the cohesin HiChIP performed with the sequential (FA-EGS) cross-linking protocol improves the signal-to-noise ratio and ChIP efficiency in comparison to the standard one (25) and enhances the detection of chromatin loops and stripes in human lymphoblastoid cells. We have developed nf-HiChIP pipeline that combines the analytical approach designed for ChIP-seq data processing (mapping, filtering, peak calling, coverage tracks calculations) with HiChIP-specific analysis (MAPS pipeline, (31)) to facilitate the user the comprehensive and timely analysis of many HiChIP datasets in parallel, without the need to perform additional ChIP-seq experiments. Furthermore we show that the data derived from the improved cohesin HiChIP protocol can be combined with the recently developed energy-based modelling approach LoopSage (35) to generate accurate biophysical models of the loop extrusion process.

**Figure 4.**
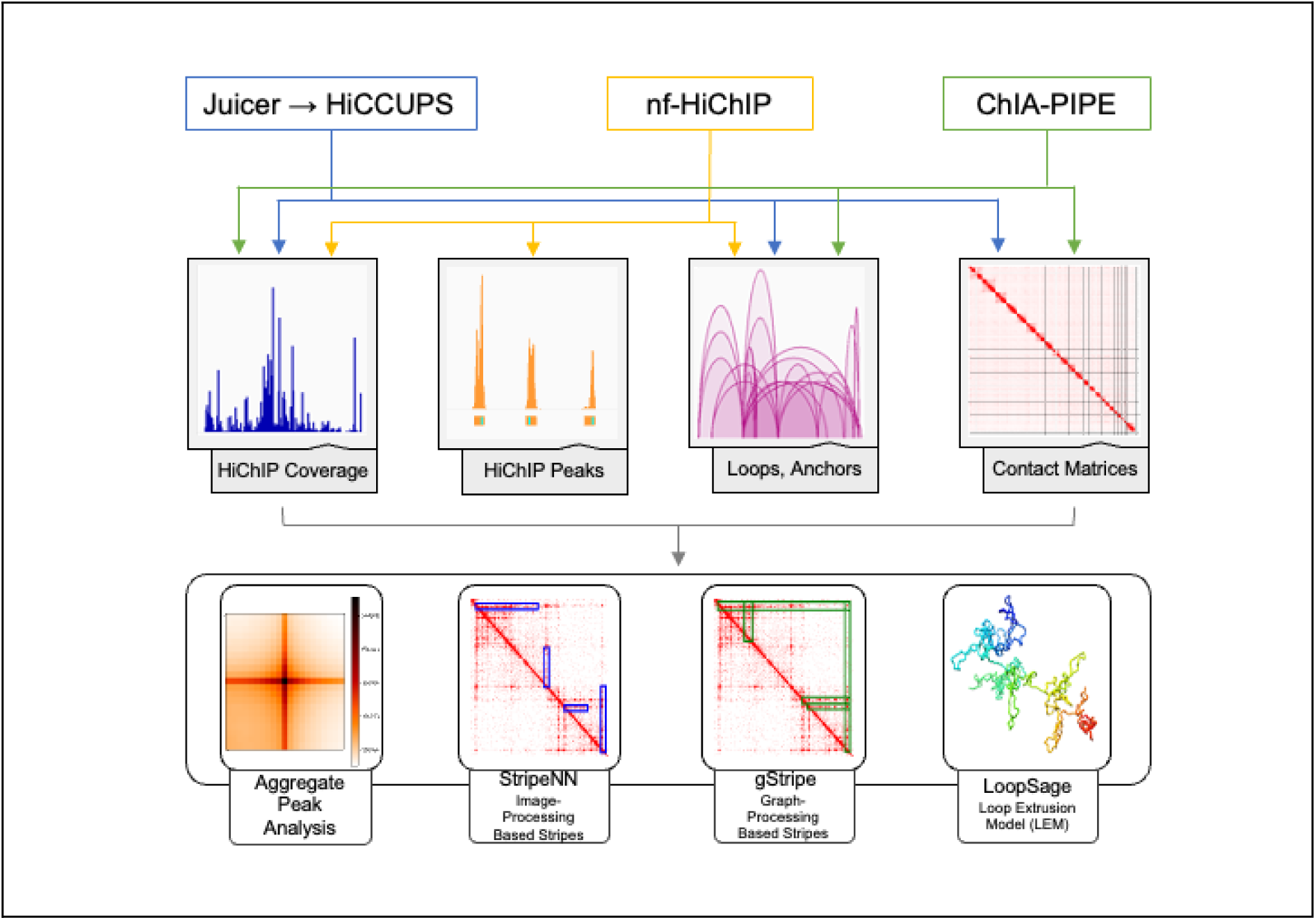
Systematic representation of the bioinformatic analyses and pipelines used in this study.

Dual cross-linking was previously reported to improve the signal-to-noise ratio and increase the sensitivity of theChIP assay in the case of proteins with hyperdynamic exchange with DNA or indirectly DNA-bound proteins (40, 41, 54). Cohesin also exhibits a dynamic association with DNA. We report that dual cross-linking improves the quality of cohesin HiChIP, which subsequently results in higher sensitivity of the experiment compared to standard cohesin HiChIP. Importantly, our results suggest that the increased efficiency of the 3C step of the protocol may be another factor contributing to the improved detection of 3D chromatin features such as loops and stripes. The use of dual cross-linking (EGS or DGS and FA) has been shown to reduce the experimental noise by decreasing the number of random Hi-C ligation events, leading to the improved loops detection (44, 55). Similarly, we observed more looping interactions in FA-EGS HiChIP, compared to cohesin FA HiChIP and these interactions were stronger. Importantly, the use of the FA-EGS HiChIP method allows the detection of 2-3 times more loops in human cells with 4-6.5 less sequencing depth compared to dual cross-linking Hi-C (44), indicating that the former method may be a more suitable and economical choice for studies focusing on the 3D chromatin structure at the resolution of loops. However, it is important to note that the influence of the cross-linking procedure on the quality of cohesin HiChIP quality might be cell type specific. For example, in the case of the undifferentiated cell line, human embryonic stem cells (56), a high signal-to-noise was obtained for the cohesin HiChIP experiment was obtained despite the use of only one cross-linking agent (FA) for fixation.

Our aim was to thoroughly investigate the results that can be obtained from HiChIP sample processing. Commonly used HiChIP data analysis tools like Juicer, HiC-Pro, and Fit-Hi-C, which were not specifically designed for HiChIP, may use normalisation and loop calling methods appropriate for Hi-C datasets. These methods may overlook the specific interactions in HiChIP. Unlike Hi-C, where a sparse interaction matrix is undesirable and is corrected using techniques such as Knight-Ruiz (KR) normalisation (6, 30, 57) to account for experimental bias, HiChIP interactions typically yield a sparser matrix. Using HiChIP-specific tools such as MAPS (31), we identified interactions that are undetectable when using KR-normalised interaction maps for loop calling. We also experimented with replacing KR-normalised maps with Vanilla Coverage (VC) normalisation (6) in the Juicer pipeline. Integrating VC normalisation, which unlike KR, does not assume a uniform distribution of interaction frequencies and thus preserves unique HiChIP-specific interactions, improved the consistency of our results. However, image-based loop calling tools such as HiCCUPS (30) may exclude closely positioned loops or those that start or end within the same matrix pixel, potentially indicating loop extrusion processes, that are important in the context of cohesin HiChIP data. These loops however, could be captured by direct processing of interaction pairs in HiChIP-focused approaches such as MAPS.

Our nf-HiChIP pipeline includes ChIP-seq-specific steps, including (1) a mapping approach that differs from the HiC mapping approach, (2) coverage track generation, and (3) peak calling that can be applied to both HiChIP and corresponding ChIP-seq samples, allowing dual analysis of HiChIP samples – both as ChIP-seq and HiChIP datasets. This dual approach provides more comprehensive outputs: peaks, coverage, and loops, from a single sample, increasing the data richness compared to traditional methods that require separate, manual analyses. This adaptability enhances the personalisation of analyses based on the experimental data available.

Importantly, nf-HiChIP is scalable and automated. Unlike conventional tools nf-HiChIP can process multiple samples simultaneously, making efficient use of available computational resources. The pipeline automates the processing of individual and pooled biological replicates, and offers extensive customization options. Users can add personalised steps, select different peak calling tools, or adjust filtering processes, to meet the specific needs of their research. The integration of Nextflow (34) and Docker (58) technologies facilitates computational tasks on clusters and could be extended to analysis on cloud services such as AWS. In the future, we plan to extend the capabilities of nf-HiChIP by including more sophisticated post-processing features, such as correlation analysis between different samples and replicates, and automated generation of statistical plots. These enhancements will further streamline the workflow and provide deeper insights into the dynamics of chromatin interactions.

The visibility of architectural stripes depends on the characteristics and quality of the heatmaps obtained. In FA-EGS cohesin HiChIP the heatmap view is sharper and has less noise in comparison to cohesin FA HiChIP. Consequently, in FA-EGS HiChIP we can see finer arrangements of stripes and chains of distinct interactions in locations where standard HiChIP shows a smooth signal. These patterns can be interpreted as obstacles in the extrusion process (intermediate states of the looping process) or as a finer arrangement of loops. Similar stripes appearance is also seen in deeply sequenced Micro-C data (55, 59), but not in standard Hi-C data, where stripes are more uniform (16, 17). Therefore, we believe that cohesin dcHiChIP provides more detailed dynamics of the loop extrusion process and a more complete picture of the interaction structure than the standard one. This allows the gStripe algorithm, which benefits from stronger and more numerous loops, to detect more stripes than the classical approach exemplified by the Stripenn tool, for which targets a smooth signal (33). Overall, the greater detail of the FA-EGS heatmap data improves the distinguishability of finer stripes in the cohesin HiChIP experiment. The detection of stripes is important, as beside its relevance for loop extrusion and super-enhancer activity, they can be associated with certain pathological phenomena, such as incorrect expression of genes (60) and increased topoisomerase activity (17), which can potentially promote cancer.

We believe that our findings will be a valuable resource for researchers who encounter difficulties in their ChIP-based and high-throughput 3C-based experiments and who are looking for tools to improve and complete their bioinformatic analysis of chromatin spatial organisation data. Importantly, the use of LoopSage 3D modelling to visualise a region of interest ensures biological realism by representing combinations of conformations that can occur in individual cells. Our approach can support studies that focus on the relationship between chromatin structure and the processes that occur in the nucleus such as DNA replication, transcription or DNA repair in which cohesin is an important player.

## Materials and Methods

### Cell line and culture conditions

Human HG00731 lymphoblastoid cell line, purchased from Coriell Institute for Medical Research, was grown in RPMI 1640 (Biowest) with 15% foetal bovine albumin (Biowest) and 2mM L-glutamine (Gibco). Cells were cultured at 37 ℃ in a humid environment containing 5% CO₂.

### Cell fixation

Cross-linking was performed using cells grown to a density of around 0.8 millions per 1 mL of culture medium at a volume of 1 ml of cross-linking solution per 1 million of cells. **FA cross-linking**. Pelleted cells were resuspended in a pre-warmed (37 ℃) RPMI1640 (Biowest) with 1% formaldehyde (Sigma, 252549) and incubated for 10 minutes with agitation. Next glycine was added to a final 0.2M concentration to quench the cross-linking reaction and the samples were incubated for 5 min with agitation at room temperature (RT). Fixed cells were then washed twice in DPBS, snap frozen in liquid nitrogen and cell pellets were stored at –80℃. **FA-EGS cross-linking.** Pelleted cells were resuspended in a pre-warmed (37℃) RPMI 1640 (Biowest) with 1% formaldehyde and incubated for 15 minutes with agitation (RT). Next glycine was added to a final 0.2M concentration to quench the cross-linking reaction and incubated for 5 min with agitation (RT). After washing once in DPBS, cells were resuspended in pre-warmed (37℃) DPBS with 2mM EGS [ethylene glycol bis(succinimidyl succinate), Thermo Scientific] agitated for 45 min at RT, quenched with 0.2M glycine and incubated with agitation for 5 min FA-EGS-fixed cells were then washed twice in DPBS, snap frozen in liquid nitrogen and cell pellets were stored at –80℃.

### HiChIP

HiChIP assay was performed according to previously described protocol (Mumbach et al., 2016) with modifications. The combination of chromatin immunoprecipitation step with HiChIP library preparation was carried out based on ChiPmentation protocol (61) as recently reported (62). The 10 million of crosslinked cells were resuspended in ice-cold 1.5 ml of Hi-C lysis buffer (10mM Tris-HCl pH 8.0, 10mM NaCl, 0.2% NP-40) supplemented with 1x <u>p</u>rotease inhibitors (PI; Roche, 04693116001), incubated on ice for 20 min and centrifuged at 1200 rpm, 5 min, 4°C. After discarding the supernatant the pellet was resuspended in 1.5 ml Hi-C lysis buffer with 1 x PI, centrifuged at 1200 rpm, 5 min, 4°C. Then supernatant was removed and the cell pellet was resuspended in 150ul of pre-warmed (65 ℃). After 5 min incubation at 65℃, 435 ul of water and 75 μl of 10% Triton X-100 (Promega HP142) was added and 15 min (37°C) incubation was carried out. Next, 75 μl of NEBuffer™ DpnII (NEB, B0543S) and 600U of DpnII (R0543M) were added to the sample and incubated for 1h at 37°C. To inactivate the restriction enzyme, the reaction was incubated at 65°C for 5 min and then cooled down to RT. To perform end repair and biotinylation the following master mix was added: 67.5 μl of water, 4.5 μl of 10mM dTTP (NEB, N0443S), 4.5 μl of 10mM dATP (NEB, N0440S), 4.5 μl of 10mM dGTP (NEB, N0442S), 45 μl of 1mM biotin-16-dCTP (Jena Bioscience NU-809-BIO16-S), 24μl of 5U/μl DNA polymerase I Large (Klenow; NEB, M0210L) and the samples were incubated for 1h at 37°C with rotation, Ligation was performed by addition to the sample of Ligation master mix (2007 μl of water, 300 μl of 10% Triton X-100, 18 μl of 10 mg/ml BSA (NEB, B9000S), 360 μl 10x T4 DNA ligase buffer (NEB, B0202), 15 μl of 400 U/μl T4 DNA ligase (NEB, M0202L), followed by 4h incubation at RT with gentle rocking. Samples were then centrifuged for 5 min at 1200 rpm, at 4°C. The nuclei pellet was resuspended in 1 ml of Nuclear Lysis buffer (50 mM Tris-HCl pH 8.0, 10 mM EDTA, 1%SDS, 1 x PI) and incubated on ice for 15 min. Sonication was carried out on Covaris S220 device in 1 ml miliTube (Covaris, 520135) using the following settings: peak power=140, duty factor= 5, cycle burst=200, time=120s, temp.= 4-8°C. After DNA shearing, the chromatin was transferred to a new tube and 100 μl of 10% Triton-X was added. To remove cell debris, samples were centrifuged for 20 min, at 4°C, at 16000xg and supernatant was transferred to a new tube. Next, the chromatin was pre-cleared by incubating with Dynabeads Protein A (Invitrogen, 10002D) coated with 10 μg of IgG (Abcam, AB171870) for 30 min at 4°C, rotating. The chromatin immunoprecipitation was carried out by overnight incubation of pre-cleared chromatin with Dynabeads Protein A coated with 10 μg of SMC1 (Bethyl Laboratories, A300-055A) or 10 μg CTCF (Abclonal, A1133) antibody at 4°C with rotation. Supernatant was then removed and the beads were washed: twice with 1 ml of Low Salt Wash Buffer (0.1% SDS, 1% Triton-X-100, 2mM EDTA, 20mM Tris-HCl pH 8.0, 150 mM NaCl), once with 1 ml of High Salt Wash Buffer (0.1% SDS, 1% Triton-X-100, 2mM EDTA, 20mM Tris-HCl p.H 8.0, 500 mM NaCl) and twice with 1 ml of 10 mM Tris pH 8.0. After careful removal of any residual 10mM Tris pH 8.0, the tagmentation reaction was carried out by resuspending the beads in tagmentation mix (15 μl 2xTD buffer, 14 μl water), adding of 1 μl of Tn5 enzyme (TTE Mix V50, Vazyme, TD501) and incubation of the reaction for 10 min at 37°C, with 530 rpm shaking, Next tagmentation reaction was removed and the beads were washed subsequently with 1 ml of Low Salt Wash Buffer, 1 ml of High Salt Wash Buffer, 1 ml of LiCl Wash Buffer (10 mM Tris-HCl pH 8.0, 250 mM LiCl, 1% NP-40, 1% Sodium Deoxycholate, 1m mM EDTA) and 1 ml of 10 mM Tris pH 8.0. To elute the DNA. The beads were then resuspended in 400 ul of freshly prepared DNA Elution Buffer (50mM NaHCO3, 1% SDS), incubated for 15 min at 65°C. The supernatant was then transferred to a new tube and reverse crosslinking and RNA digestion was performed by adding 44 μl of mix consisting of: 20μl 5M NaCl, 8 μl 0.5 EDTA, 16 μl Tris 1M pH 8.0 as well as 8 μl of 10 mg/ml RNAse (Thermo Scientific, EN0531) and incubation for 6h at 65°C. Proteins were then digested by 1h incubation at 55°C with 4 μl of 20 mg/ml Proteinase K (Ambion, AM2546). DNA was then recovered by phenol/chloroform extraction and ethanol precipitation. The DNA pellet was resuspended in 21 μl of 10mM Tris pH 8.0, the concentration was measured with Quantus Quantifluor ONE dsDNA system (Promega, E4870) and the sample volume was increased up to 100 μl with 10mM Tris pH 8.0. Next, 10 μl of MyOne Streptavidin C1 Beads (Invitrogen, 65001) were washed twice with 400 μl of Tween Wash Buffer (5mM Tris-HCL pH 7.5, 0.5mM EDTA, 1M NaCl, 0.05 % Tween-20), resuspended in 100 μl of 2x Biotin Binding Buffer (10mM Tris-Hcl pH 7.5, 1MM EDTA, 2M NaCl), mixed with the DNA sample and incubated for 20 min with rotation at RT. The beads were then washed (1) twice with 600 μl of Tween Wash Buffer, (2) once with 100 μl 10mM Tris pH 8.0, (3) once with 50 μl 10mM Tris pH 8.0 and resuspended in 23 μl of 10mM Tris pH 8.0. To prepare sequencing libraries on-bead PCR amplification was performed with a total of 12 PCR cycles using Truseq indexes. The libraries were then sequenced on Illumina NovaSeq 6000 platform at the Genomics Core Facility, CeNT, University of Warsaw.

## Data Processing and Analysis

### nf-HiChIP pipeline

Both HiChIP and ChIP-seq datasets were processed using the nf-HiChIP pipeline. The initial stage of the pipeline mirrors a ChIP-seq analysis framework. Sequencing reads were first aligned to the hg38 reference genome employing the BWA-MEM algorithm (version v0.7.17) (63). Subsequently, SAMtools (64) was utilised to filter the aligned reads, retaining only those with a mapping quality (MapQ) score greater than 30. This step was followed by deduplication of reads to eliminate potential biases arising from PCR amplification. BAM files were converted to normalised read coverage tracks (bigwig format) using the program bamCoverage in the deepTools package (45) with the binSize parameter set to default and using the reads per kilobase per million mapped reads (RPKM) normalisation option. Coverage tracks were generated using the deepTools package, facilitating the visualisation and analysis of read depth across genomic regions.

Peak calling was performed with the MACS3 software (version v3.0.1) (65) adopting a stringent q-value cutoff of 0.01 to identify regions of significant enrichment. The outcomes of peak calling, applied to both ChIP-seq and HiChIP data, served as input for the MAPS algorithm (31), which delineates the final list of chromatin interactions with the anchor width of 5kb from each sample. This analysis was conducted individually for each replicate before pooling data to create a comprehensive dataset for each sample, whereupon the entire analysis was reiterated to ensure robustness and accuracy.

Final outputs generated by the pipeline encompass a range of files for each replicate and sample, including: (i) coverage tracks, (ii) identified peaks, and (iii) comprehensive MAPS output detailing the called pairwise interactions. This structured approach ensures a systematic and reproducible analysis pipeline, facilitating the in-depth exploration of chromatin dynamics and interactions. All the intermediate files (reads mapped to the reference genome, indexed BAM files, coverage tracks) can be easily accessed too.

### Juicer

Furthermore, all HiChIP samples were processed using the Juicer pipeline (Version 1.22.01) (30), a standard for Hi-C and HiChIP data analysis. Notably, the commonly used Knight-Ruiz normalisation (KRnorm) for Hi-C data was found inadequate for HiChIP datasets, often obscuring characteristic interactions and stripes. Consequently, VCnorm normalisation was employed with Juicer’s HiCCUPS for interaction calling, circumventing the aforementioned issues.

Aggregate peak analysis (APA) was conducted to assess the quality of loop calling by nf-HiChIP, HiCCUPS and ChIA-PIPE across samples. This analysis, performed via the Juicer apa function, was visualised using Matplotlib, providing a detailed comparison of loop calling efficacy. As recommended in (6) for each plot, we used the P2LL as APA score to determine enrichment of the signal to the lower left corners.

### HiChIP quality analysis and data visualisation

Post-analysis involved comprehensive sample comparisons through custom scripts in Jupyter notebook (66), utilising Numpy (67) and Pandas (68) for basic analysis, with Matplotlib and matplotlib_venn for visualisations. Sample correlation, peak enrichment, and fingerprint analysis employed deepTools (69), while genomic tracks and interaction maps were visualised using IGV (70) and Juicebox (71), respectively.

The Pearson correlation was computed for the coverage files using the multiBigwigSummary module of deeptools (69), employing default parameters to determine the average scores across various genomic regions from the provided set of bigwig files. The resulting data was then utilised by the plotCorrelation module to generate correlation coefficients, which were subsequently visualised using Matplotlib as a clustered heatmap. In this representation, the colours signify the correlation coefficients, while the clusters are formed using complete linkage.

We have implemented custom in-house scripts to pinpoint overlaps among significant interactions, stripes, and stripe domains between all HiChIP samples. To estimate the overlap of loops between the samples, we set a 15kb tolerance from the length of both anchors. Since one-to-many overlaps are possible for two sets of regions (anchors, stripes, etc.), the resulting matrices are not symmetric. For two stripes to be considered overlapping we require that they have the same orientation (horizontal or vertical) and their anchors intersect with 10kb tolerance and we do not consider the lengths of the stripes.

### ChIA-PIPE

The ChIA-PIPE (32) pipeline was used to process and map the CTCF HiChIP and Cohesin HiChIP data to the human hg38 reference genome. First, the reads were looked for the ligated restriction enzyme site (called a pseudo linker in HiChIP), and only the sequences that had the pseudo linker were kept for linker trimming. The flanking sequences were then mapped to the human reference genome (hg38) by Burrows Wheeler alignment (BWA), and only uniquely aligned sequences (MAPQ ≥ 30) were kept for deduplication. Next, reads with a linker sequence detected with both ends having genomic tags are used for the detection of interaction loops. These reads are categorised as self-ligation PET (both ends of the same DNA fragment) or inter-ligation PET (both ends from two different DNA fragments in the same chromatin complex). Inter-ligation PETs with a genomic span of ≥8 kb are illustrated to represent the long-range interactions of interest, further subdivided into intra– and inter-chromosomal PET clusters. Lastly, different PET clusters in the same protein factor’s binding peak region were joined together with 500-bp extensions to make merged anchors. Only loops with both anchors supported by peaks identified by nf-HiChIP pipeline were retained for each sample. Only clustered and merged intra-chromosomal PETs with a PET count ≥ 3 and a genomic span < 1MB were retained as reflecting the chromatin interactions of interest.

### CTCF motifs

To obtain CTCF motifs, we have taken a two-way approach. The first one is plainly using the motifs module from Biopython. We use it to obtain, e.g. anchors that have motifs (with a certain score – usually 7.0) in their sequence. However, to produce more detailed models, we have also introduced probabilistic motifs. In that case, we are taking all the signals of motif presence available, and we are calculating the probabilityof the motif being upstream or downstream-oriented using the following formula:

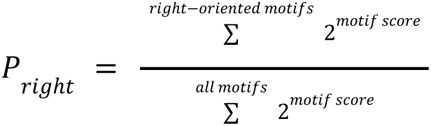

In the equation, we are taking all scores (that are in log2 form) of all motifs present in the sequence, and we convert them back from log2 form to regular one. Then, we sum the scores for the right/left orientation, and divide it by all scores from both right and left-oriented motifs. That way, we can say, e.g. that the probabilistic score of the motif being left is 30%, and right is 70%. That approach is useful for modelling and simulations.

### Calling architectural stripes

Stripe calling was performed using two tools: the Stripenn (33), which operates on heatmap data, and gStripe, which was developed for calling stripes from sets of discrete interactions, such as loops obtained from other tools. The Stripenn is based on image analysis and computer vision methods. It processes the contact matrix obtained from a 3C experiment using a series of computer vision techniques, most importantly the canny edge detection, to select regions with high “stripeness” scores, i.e. those that are visually consistent with a horizontal or vertical stripe. It also calculates the p-value of a stripe, based on the distribution of the values taken from the same heatmap, and this provides the final significant stripe set. To ensure our analyses reflect the best possible extraction of useful information from each dataset, we tuned two of Stripenn’s parameters (the Canny edge detection parameter and the kernel size of the mean filter) to obtain the maximum number of stripes in each dataset. This was done by a simple grid search centred around the values suggested by the authors (see **Supplementary Table 8**).

As an alternative to image processing methods, we also include the results from our own tool, gStripe, designed specifically for sparse data. In contrast to contact matrix-based methods, gStripe operates on sets of loops obtained from other tools, in our case nf-HiChIP, HiCCUPS and ChIA-PIPE. The algorithm can be divided into three parts: preprocessing of input data, identification of candidate stripes, and selection of final stripes. During preprocessing all input loop anchor coordinates are binned at 5 kb resolution, to avoid any bias introduced by comparison between binned and non-binned input data. Next, a graph representation of the chromatin interactions is created. Continuous clusters of overlapping anchors are merged to represent a single vertex in the graph. The edges are then added to the graph, where there are interactions between anchors assigned to two different vertices. Multiple edges connecting the same vertex are collapsed into a single edge, with weight equal to the number of interactions (hence, the resulting graph is not a multigraph). The physical position of each vertex is calculated as the centre of the anchor cluster forming the vertex.

To identify the candidate stripes, each vertex with at least one downstream neighbour in the graph is selected as a potential anchor of a horizontal stripe, i.e. a stripe which appears as a horizontal flare above the diagonal on a heatmap, or whose anchor (the location where the stripe reaches the diagonal) precedes its other end in terms of genomic coordinates. Likewise, each vertex with at least one upstream neighbour is selected as a potential vertical stripe (appearing as vertical pattern above diagonal on a heatmap, with the anchor being downstream of the stripes longest extent). We will now refer to these anchor-forming vertices simply as “anchors”, and to the other vertices in a candidate stripe as “leaves”. A number of values is computed for each leaf v_i_, in order to measure the quality of the stripe. Let ʋ*_i_* = 1, 2, …, *k* denote the k leaves of a stripe, and v_0_ denote the anchor.

1. The length of the stripe at the leaf ʋ*_i_*: *d_i_* = |ʋ*_i_* − ʋ*_i_* |. Prior to this step, the vertices are arranged so that *d_i_* is ascending with i, i.e. ʋ_0_ is the anchor, ʋ_1_ is the vertex closest in terms of genomic coordinates to the anchor, etc. In a horizontal stripe, the coordinates of ʋ*_i_* have increasing order, and in a vertical stripe they have a decreasing order.
2. Relative gap 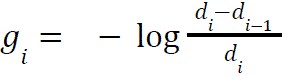. All lengths are set to be at least 1kb to ensure that this value is well-defined.
3. *q_gap*_i_ – quantile of the relative gap values across all candidate stripes in the dataset identified in the previous step.
4. *stripe_score*_i_ – quadratic mean of the *q_gap_j_* for j = 1,…, i. Represents the average quality of the stripe of up to a certain point (i.e. leaf ʋ*_i_*).
5. *cross_score_i_* – each leaf is potentially a part of two orthogonal stripes: horizontal and vertical one. For a given stripe direction the cross score of ʋ*_i_* is the ratio of *stripe_score*_i_ calculated in that direction and the stripe score in the orthogonal direction.
6. The length of the stripes are determined: a stripe is considered to end with a leaf ʋ*_i_*, if *g_i_*_+1_ drops below 0.05, or if there are no further leaves.
7. If two adjacent stripes with the same directionality overlap and differ in length by no more than 200kb, they are merged together.
8. Only stripes with at least two leafs, and having a minimum length of 20kb are selected and considered further.

The above steps result in a set of candidate stripes. In order to select the final stripes from the candidate set, two features of the stripes are considered: the *stripe_score* of the leaf terminating the stripe (note that this is a form of cumulative quality of the stripe up to this leaf), and the mean *cross_score* of every leaf in the stripe after its length has been trimmed. These parameters measure the continuity and distinctiveness of the stripe respectively, and the selection of the final stripe set is done by placing a lower threshold on these values. The choice of the thresholds is done by visual inspection of the output and manual adjustment; this does not require, however, re-running the algorithm. We used the thresholds of 0.45 for *stripe_score* and 0.9 for *cross_score*. Note that in contrast to methods based on heatmap analysis (such as Stripenn), potential stripe locations are already narrowed down, as the stripe can only be called in a location where two consecutive loops were present.

### Biophysical modelling

For loop extrusion modelling, we used LoopSage (35). We assume that the stochastic system can be described by a Boltzmann distribution, where the temperature parametrises the rebinding probability of cohesin (72). The temperature decreases during the simulation following a Simulated-Annealing approach (73). For the purposes of this work, the polymer consists of 1000 monomers with 50 cohesins that play the role of loop extruders. The stochastic simulation ran for 10^4^ Monte Carlo steps with a sampling frequency equal to 100 and a burn-in period equal to 10^3^. The folding and binding coefficients of the energy of the stochastic simulation were chosen to be equal to the default ones f=b=-1000 (35), whereas for the crossing penalty, we chose k=20000.

The simulation pipeline is composed of two main parts: a stochastic simulation part, where we compute the stochastic trajectories of cohesins in the DNA fibre by running the Monte Carlo simulation, and a molecular simulation part, where we import the cohesin positions exported by the numerical simulation part and we feed them into an OpenMM model (74–76) which is capable of producing 3D-structures from them. Having produced the ensemble of models, we compute the average inverse all-versus-all distance heatmap of each one of these 3D models. Finally, we aggregate them into an average final heatmap, which should reconstruct similar patterns as the experimental HiChIP heatmap.

## Data availability

Sequencing data generated in this study is deposited in the Gene Expression Omnibus (GEO) database, with an accession number GSE266640. nf-HiChIP pipeline is available at https://github.com/SFGLab/hichip-nf-pipeline. The pipeline is implemented in Nextflow with Docker support and processes the output of various tools. The gStripe algorithm implementation is available at https://github.com/SFGLab/gStripe as a Python 3 package. The LoopSage, an energy-based model for loop extrusion, is available as an open-source project on GitHub at https://github.com/SFGLab/LoopSage.

## Acknowledgements

We would like to thank Victor Corces and the Corces lab for the introduction to the HiChIP assay and analysis, Dorota Adamska from Genomics Core Facility (CeNT, UW) for designing the sequencing strategy and providing constant support in troubleshooting experiments, our laboratory members for discussions and Veronika Mančíková for proofreading of the manuscript.

## Author contributions

K.J., A.A., Z.P.-T., and D.P. conceived the study. K.J. performed HiChIP experiments; Data analysis and pipeline development was performed by Z.P.-T. and A.A. with assistance from M.C, K.B and D.P. M.D. performed stripe calling and analysis, and S.K. carried out loop extrusion modelling. D.P. and K.J. designed and supervised the study. K.J., A.A., and Z.P.-T prepared the manuscript with assistance from M.C., M.D., S.K. and D.P. All authors discussed the results and commented on the manuscript.

## Funding

This project received funding from the Polish National Science Centre (2019/35/O/ST6/02484 to DP, 2020/37/B/NZ2/03757 to DP, 2020/04/X/NZ2/01006 to KJ), Boehringer Ingelheim Fonds travel grant to KJ, Warsaw University of Technology within the Excellence Initiative: Research University (IDUB) programme, National Institute of Health USA 4DNucleome grant 1U54DK107967-01 “Nucleome Positioning System for Spatiotemporal Genome Organization and Regulation”, EU-funded Innovative Training Network “Molecular Basis of Human Enhanceropathies” (Enhpathy, www.enhpathy.eu), the Horizon 2020 research and innovation programme under the Marie Sklodowska-Curie grant agreement No 860002 and Excellence Initiative: Research University (IDUB) programme with Agreement no. BOB-IDUB/IV.4.1/4/2024 to AA, Polish National Agency for Academic Exchange (PPN/STA/2021/1/00087/DEC/1 to MC). Computations were performed thanks to the Laboratory of Bioinformatics and Computational Genomics, Faculty of Mathematics and Information Science, Warsaw University of Technology using Artificial Intelligence HPC platform financed by Polish Ministry of Science and Higher Education (decision no. 7054/IA/SP/2020 of 2020-08-28). NGS was performed thanks to Genomics Core Facility CeNT UW (RRID:SCR_022718), using NovaSeq 6000 platform financed by Polish Ministry of Science and Higher Education (decision no. 6817/IA/SP/2018 of 2018-04-10)

## Conflict of interest statement

None declared.

## Notes

### Competing Interest Statement

The authors have declared no competing interest.

